# Chitosan diet alters the microbiome of adult house flies

**DOI:** 10.1101/2022.08.31.502951

**Authors:** Hila Schaal, Mallory J. Choudoir, Vedang Diwanji, John Stoffolano, Kristen M. DeAngelis

**Author notes:** **Correspondence:** Kristen DeAngelis.

## Abstract

House flies are disease vectors, carrying human pathogens which include *Escherichia coli* and *Vibrio cholera*. To explore the use of chitosan as a bioinsecticide, we evaluated the effects of a chitosan-amended diet on *Musca domestica* (house fly). We first conducted longevity experiments to understand the impact of chitosan on house fly longevity. We confirmed that chitosan diet amendment is associated with reduced longevity and that this is not due to starvation. We then extracted fly microbiome DNA and used 16S ribosomal RNA gene amplicon sequencing and quantitative PCR to assess the composition and load of the microbiome for flies fed chitosan-amended diets compared to controls. Diversity of the chitosan-fed fly microbiomes was lower than the control, with significant dissimilarities in community composition. Chitosan-fed flies showed lower *Ralstonia* relative abundance but increased relative abundance of *Serratia*. Both control and chitosan-fed flies had highly uneven communities, but the control flies were dominated by genera *Ralstonia* and *Providencia*, while the chitosan-fed flies were dominated by genera *Serratia, Kosakonia*, and *Providencia*. Contrary to our expected results, chitosan-fed flies also contained 56% more bacteria compared to controls. Gut microbiome changes appear to result from chitinolytic bacteria becoming more relatively abundant, and our results suggest that chitosan-amended diet alters the house fly microbiome resulting in higher fly mortality.

## 1 Introduction

Chitosan is an industrially produced N-deacetylation product of chitin that has made its way into food, medical, and textile industries (Kong et al. 2010). Chitin, the second most common polymer in nature after cellulose, is commonly extracted from the exoskeleton of crustaceans through demineralization, deproteinization, and decolorization steps (Kong et al. 2010). After the extraction of chitin, a strong alkaline treatment or an enzyme for deacetylation of the N-acetylglucosamine polymer can be applied to produce chitosan (Hahn et al. 2020). Chitosan is used against plant pathogens, such as fungi as an antimicrobial agent (Orzali et al. 2017), against termites (Telmadarrehei et al. 2020), and drosophila (Pinero et al. 2021). Chitosan is even being explored for its human use as an anti-obesity theraputic (Pan et al. 2018). In their review, Rabea et al. (2003) noted the potential use of chitosan in the agricultural setting but added that its use as an antimicrobial chemical and its mode of action needs more research. Rabea et al. (2015) studied the toxicity efficacy of N-(40-propylbenzyl) chitosan and N-(2-nitrobenzyl) chitosan on *Ceratitis capitata* adults (Mediterranean fruit flies) and found significant toxic effects on both sexes. Other than increasing mortality (Stoffolano et al. 2020), few studies have been conducted on the effect of chitosan on *Musca domestica* (house flies).

The adult house fly is a vector of numerous pathogens of humans (Stoffolano, 2019) with costs of $1 billion per year in the U.S. (Geden et al., 2021). Most epidemics involving human pathogens are transmitted via biotic factors and disease vectors including insects (Bahrndorff et al 2017). To promote good public health, there is a need to control the widespread populations of the house fly. In addition, the house fly poses a serious food safety threat because adults can rapidly transfer human pathogens that can easily enter our food supply. The house fly is known to carry over 100 pathogens, some that can be life-threatening. Adults can potentially spread disease agents to human populations, such as *Escherichia coli, Salmonella enterica* serotype Typhi, *Shigella*, and *Vibrio cholera* (Khamesipour et al. 2018, Brewster et al. 2021). Commonly used methods of fly population control include avoiding attracting flies by not leaving food out for an extended period of time, or the usage of fly tape traps or chemicals to kill the flies (Pickens and Miller 1987). However, the maintenance of sticky traps can be an inconvenience and does not completely deter disease vectors. In a major review of the current management strategies and research needs, Geden et al. (2021) stressed two major points: screening new active ingredients for toxicity and the need to avoid the widespread resistance issue of the fly to nearly every insecticide currently on the market.

The low nutrient needs and short life cycle of the house fly, approximately 28 days, also makes them good model systems to study development and dynamics of the microbiome (De Jonge et al. 2020). Flies are commonly used as model organisms for studying the link between diet, longevity, and microbiome. The disruption of the fruit fly microbiome by bacterial infection and ingestion negatively impacts the longevity of the organism (Behar et al., 2008), while it was separately observed through gnotobiotic experiments that flies with higher gut microbiome diversity exhibit a fitness tradeoff, with more diversity resulting in flies that mature and reproduce more quickly, but also die sooner (Gould et al., 2018). Antibiotics significantly promoted the longevity of sugar-fed male and female medflies but only when they were fed nutritionally incomplete diets (Ben□Yosefet al., 2008), an effect that was also observed in fruit fly reproduction (Andongma et al., 2018). These studies illustrate the tight link between diet, microbiome, and longevity (Engel and Moran, 2013), and support diet as a target for biocontrol.

Chitosan is a bioactive, environmentally safe, inexpensive chitin derivative that has now been used in agriculture. As a novel ingredient, any resistance mechanism to chitosan will be different than those currently used against insecticides being used. A few studies have been done on two other adult flies, while a laboratory study has shown that chitosan added to a sugar bait kills adults (Stoffolano et al., 2020). In this study of flies grown under laboratory grown conditions, a sugar diet amended with chitosan effectively increased mortality in adult house flies compared to a sucrose control diet. Within 4 days of 2% chitosan-diet ingestion, mortality reached 100%, compared to the control group, where mortality reached 100% by day 13. Its mode of action is unknown, thus the focus of this research.

We tested the hypothesis that chitosan fly mortality is associated with adverse effects on the microbiome of house flies. Using the same model organism as Stoffolano et al. (2020), the control group of house flies were fed a sucrose diet while the experimental group of flies were fed a sucrose diet amended with chitosan. In repeating Stoffolano et al. (2020) survival experiment, we added a starvation group to compare and validate chitosan’s effect on fly longevity. We used bacterial 16S rRNA genes amplicon sequencing from whole fly DNA extractions to study the composition of the microbiome of the fly guts. Using ASV (amplified sequence variant) data, microbiome diversity and composition is compared between the chitosan group and the control group, which we expected to vary based on prior studies (Gupta et al. 2012, Park et al. 2019). To date, no papers have been found showing the effect of a chitosan diet on the insect microbiome. We expect that the chitosan fed flies will have a decrease in the gut microbial diversity and abundance, therefore affecting longevity. Understanding the relationship between chitosan and the house fly microbiome may lead to a revolutionary pesticide solution for house fly population control. Prior to its general acceptance as a commonly used pesticide, there needs to be a better understanding of the antimicrobial effects of chitosan as a tool for insect population control for farms and public health.

## 2 Materials and Methods

### 2.1 Preparing and feeding house flies

House fly pupae were sourced from Dr. Christopher Geden located in Gainesville, Florida. Three separate sterilized cages were set-up for the three different groups: control, chitosan-fed, and starvation male and female flies. To prepare for the fly experiments, cages were cleaned using dish soap and water, air dried for 1 hour, then sprayed with 70% ethanol. All cages contained a sealed container filled with 20 mL water, where a moist single Absorbal® dental wick was threaded through two punched holes on the lid. In the control cage, the food source contained sugar water as 10% sucrose, with 2% ascorbic acid added to help dissolve the chitosan in the experimental diet. In the chitosan cage, the food source contained sugar water with dissolved chitosan powder (10% sucrose, 2% ascorbic acid, 2% chitosan powder). In the starvation cage, water was given, but there was no food source given. The chitosan diet was prepared with low molecular weight chitosan (Sigma Aldrich®, 75%-85% deacetylated 50-190 kDa). We scooped approximately one teaspoon of fly pupae into each cage, or about 5 mL containing an estimated 103 pupae. One day after the pupae emerged into adult flies, they would have their first feeding on their specified food. Three days after the first feeding, adult flies would be euthanized by freezing them for approximately 15 minutes at - 20 °C. The sexes of house flies can be differentiated by morphology. Female flies were bigger in size, and males’ eyes holoptic while females’ eyes are dichoptic. We used live flies for the microbiome analyses, separate from the longevity feeding experiments, though they were set up in the same manner. To measure the microbiome of live flies, they were euthanized via freezing at day 3 after the first day of eating.

### 2.2 Whole Fly DNA Extraction

To measure the effect of chitosan on gut microbiome communities, we pooled and extracted DNA from whole flies. The forceps and the surrounding environment were disinfected with 70% ethanol before, during, and after pooling. An ethanol burner was used to heat and sanitize the forceps before the experiment. To cool down the tools, they were dipped in 70% ethanol before contacting the flies. 1.5 mL tubes were prepared according to the following factors: control male (CM), control female (CF), chitosan male (HM), chitosan female (HF). Flies were externally sterilized by dipping them in 70% ethanol for approximately 30-60 seconds, then placed in a 1.5 mL tube containing 150 μL PBS (phosphate-buffer saline). Exactly three flies of each group were pooled into a single tube to achieve approximately 50 mg of tissue. Four replicates (tubes) of flies were included from each treatment group.

Following the Qiagen DNA Blood and Tissue kit protocol, total DNA was extracted from house flies according to the kits’ insect supplement guide (Qiagen, Germantown, MD). The manufacturer’s protocols were followed with the exception of using 100 μL of buffer for the elution step to extract a higher DNA concentration from our samples. The DNA concentration was measured via Invitrogen™ Qubit™ DNA Assays (Thermo Fischer Scientific, Waltham, MA).

### 2.3 Quantitative Polymerase Chain Reaction (QPCR) of bacteria

To detect variations in bacterial load between control and chitosan house fly microbiomes, we quantified the bacteria from extracted DNA using 16S ribosomal RNA (rRNA) gene universal primers targeting the V4 hypervariable region 515f (5’-GTGYCAGCMGCCGCGGTAA-3’) (Parada et al. 2016) and 806r (5’-GGACTACNVGGGTWTCTAAT-3’) (Apprill et al. 2015). The thermocycler program consisted of an initial denaturation step at 95°C for 3 min followed by 40 cycles of 90°C for 30 s, 50°C for 30 s, 72°C for 30 s and a final extension at 72°C for 10 min. Copies of 16S rRNA gene were determined based on a standard curve constructed from amplicons generated from a lab bacterial strain *Agromonas* sp. GAS125 template and quantified using Qubit and Nanodrop. Standard curves with an R^2^ of 95 or higher were considered to have a strong level of detection (Kralik and Ricchi, 2017). Based on suggestions from New England BioLabs (Ipswich, MA) for a sample size of 50 μL, approximately 50-250 ng of DNA is required for a detectable read. We used 20 ng DNA for QPCR with a sample size of 20μL.

### 2.4 Gene Sequencing for Community Analysis

We amplified and sequenced the V4 hypervariable region of the 16S rRNA gene using the same 515f/806r barcoded primers as previously described (Caporaso et al. 2011; Parada et al. 2015) following the standard Earth Microbiome Project 16S Illumina amplicon protocol (Caporaso et al. 2018). PCR amplification was performed in duplicate for 16 whole fly extractions alongside a DNA extraction negative control and a no template (PCR) negative control. Each 25 μl reaction contained 10 μl Invitrogen Platinum Hotstart Master Mix (Thermo Fisher Scientific, Waltham, MA), 13 μl PCR-grade water, 0.5 μl forward and reverse primers (10 μM), and 1 μl DNA template. The thermocycler program consisted of an initial denaturation step at 94°C for 3 min followed by 35 cycles of 94°C for 45 s, 50°C for 60 s, 72°C for 90 s and a final extension at 72°C for 10 min. Duplicate amplification samples were pooled, cleaned, and normalized using a SequalPrep Normalization plate (Thermo Fisher Scientific, Waltham, MA). Amplicons were sequenced at the University of Massachusetts Amherst Genomics Resource Lab (Amherst, MA) on the Illumina Miseq instrument using 2×150 bp paired-end read chemistry. Samples were pooled with samples from another project and sequenced on a single run.

Amplicon sequence variants (ASVs) were inferred using the DADA2 (v3.10) bioinformatic pipeline (Callahan et al. 2016). Raw sequences were demultiplexed and primer sequences were removed using cutadapt v1.9.1 (Martin, 2011). Forward and reverse reads were filtered and trimmed using the following parameters: truncLen=c(140,150), maxN=0, maxEE=2, truncQ=10, rm.phix=TRUE. Sequence inference was performed using error rates determined from quality filtered sequences, paired reads were merged, and chimeric sequences were removed. ASV taxonomy was assigned using the DADA2 naïve Bayesian classifier method trained on the SILVA reference database v138 (McLaren, 2020). Prior to downstream analysis, we removed sequences from taxa that were present in the DNA extraction and no template (PCR) negative controls, any sequences that contained Ns, and sequences that did not resemble 16S rRNA gene sequences. Samples were rarified to normalize to relative abundance using phyloseq (McMurdie & Holmes, 2013) for all downstream analyses. These ASV sequences are available in GenBank under accession numbers OL416221 - OL416327.

### 2.5 Statistical analysis

We used t-tests to evaluate statistical differences in microbial quantities from qPCR data between chitosan-amended and control diet treatments, as well as between male and female flies. We evaluated differences based on a P-value cutoff of <0.05. We analyzed the contribution of diet and sex to the structure of the house fly bacterial communities via Principal Coordinate Analysis (PCoA) and Non-metric Multidimensional Scaling (NMDS) analysis, with dissimilarity matrices for ordinations computed using Unifrac and Bray Curtis distances for PCoA and NMDS, respectively. We analyzed the community structure using both PCoA and NMDS because we have no way of knowing whether the distance relationships are linear (as is assumed for PCoA but not NMDS). These two models have different distance matrices and assumptions, so trends observed using both models are more robust and more likely to reflect true trends. We used permutational ANOVA (R package vegan) to test if for variation in microbiomes explained by sample variables, with statistical significance evaluated following 999 permutations. All data was analyzed using R version 4.1.0 (R Core Team, 2013), and packages vegan (Oksanen et al., 2007), phyloseq (McMurdie & Holmes, 2013).

## 3 Results and Discussion

### 3.1 Starvation and Chitosan Diets Vary Longevity of House Flies

We performed two longevity trials. In the first longevity trial, the starvation group (n=100) adult flies reached 100% mortality within day 5. The chitosan group (n=150) adult flies reached 100% mortality within 7 days, as compared to the control group (n=135) adult flies that reached 100% mortality within 11 days (**Figure 1**). In the second longevity trial, the starvation group (n=60) adult flies reached 100% mortality within day 5. The chitosan group (n=80) adult flies reached 100% mortality within 7 days. Due to disruptions related to the COVID-19 pandemic, we were unable to monitor full mortality in the control group before public health restrictions were applied towards laboratory personnel. However, by day 11 approximately 75% of the control group reached mortality (**Figure 1**).

**Figure 1.**
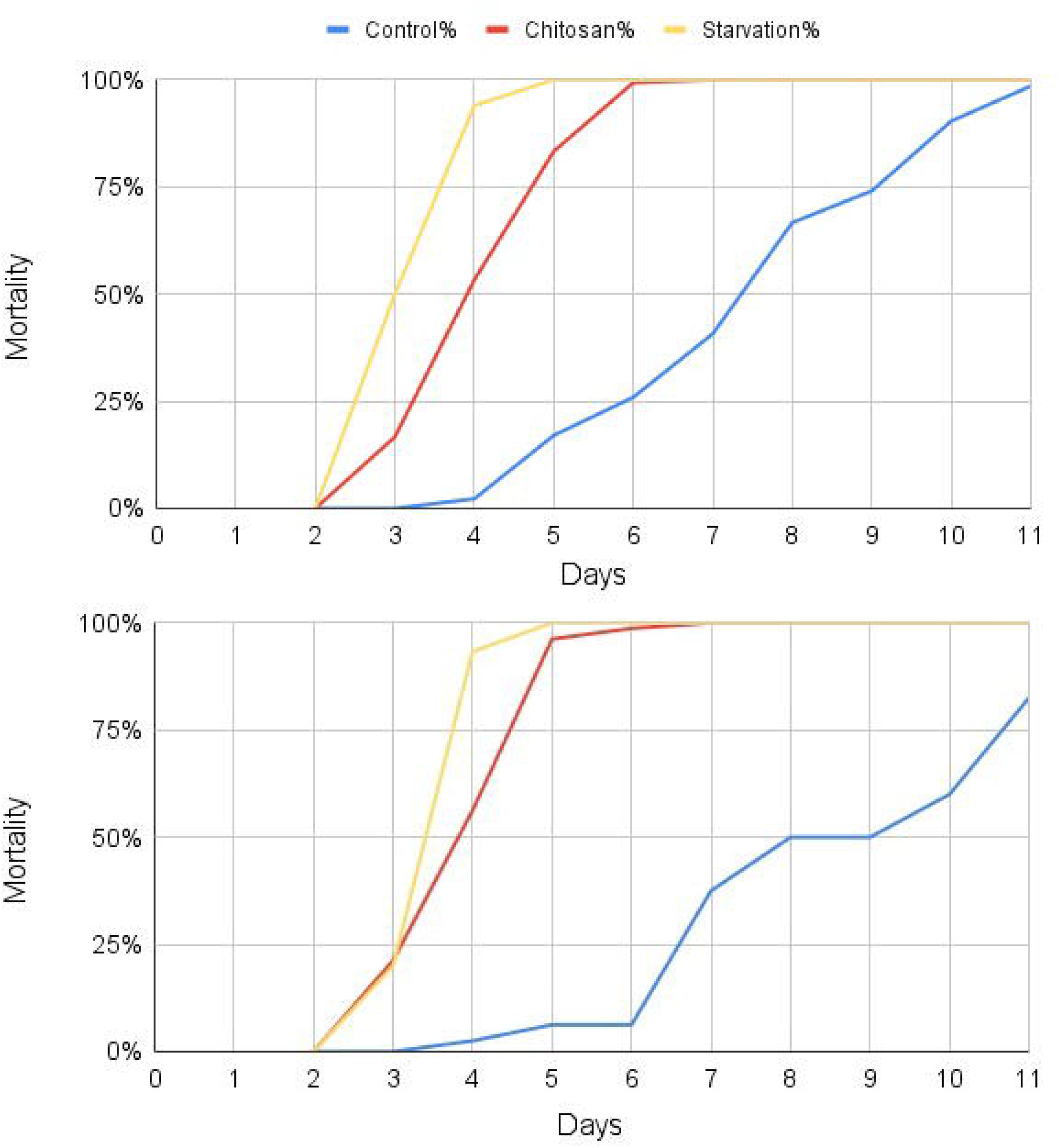
(A) Control N=135, Chitosan N=150, Starvation N=100. By day 5, 100% of the starvation group died. By day 7, 100% of the chitosan group died. By day 11, 100% of the control group died. (B) Control N=80, Chitosan N=80, Starvation N=60. By day 5, 100% of the starvation group died. By day 7, 100% of the chitosan group died. Between day 6 and 7 there was a large control group fly mortality, not due to diet, but due to flies escaping or drowning. Day 11 was the last day of data collection due to disruptions related to the COVID-19 2020 pandemic.

Our observed rate of mortality for flies fed with chitosan (100% by day 7) was slower than that observed previously (100% by day 4, Stoffolano et al. 2020), and there were a few minor caveats to our experimental approach for the mortality trials that could explain this. When we set-up the liquid diets, we put lids on the cups and string a wick through a hole in the lid for the flies to access the diet. However, some flies went through the hole and unfortunately died from drowning. This skewed some of our results due to the population counts decreasing for reasons other than diet. Some of the cages may have had small holes that flies were able to escape from, so missing flies might have been counted as dead. These issues were not specific to any one treatment type. Our replication of previous results suggests that these caveats only affect the absolute rate of mortality and not the relative effect of chitosan-amended diet on mortality.

Observationally, we noticed a behavioral difference in reactions between the fly groups. When we bumped our hands against the cage, the control flies flew all around the cage reacting to the noise. The chitosan group seemed to lack energy and had slowed movements. The starvation group had no reaction at all. At times we used forceps to poke at flies’ feet of the presumably dead flies, and sometimes, we would see a reaction showing they were still alive. This was an indication that fly health was quantitatively degraded in addition to quantified increased mortality.

### 3.2 Chitosan alters gut community composition

To test our hypothesis that a chitosan diet reduces fly microbiome alpha diversity and alters community structure, we surveyed bacterial 16S rRNA gene sequences of extracted whole fly DNA. Amplicon sequencing yielded a total of 107 ASVs, with a range of 3,467 to 14,548 sequences per sample after the quality control.

Based on PCoA and NMDS ordination models, diet strongly altered community structure (**Figure 2**). Community structure varied significantly by diet (NMDS PERMANOVA, F= 28.831, P<0.001) and explains 68% of the variance in the data (based on the model R-squared). Sex does not influence community structure (PERMANOVA, F=1.084, P<0.274 n.s.) and explains 2.6% of the variance in the data (based on the model R-squared). Consistent with previous studies (Behar et al. 2008), sex does not contribute to differences in the microbiome.

**Figure 2.**
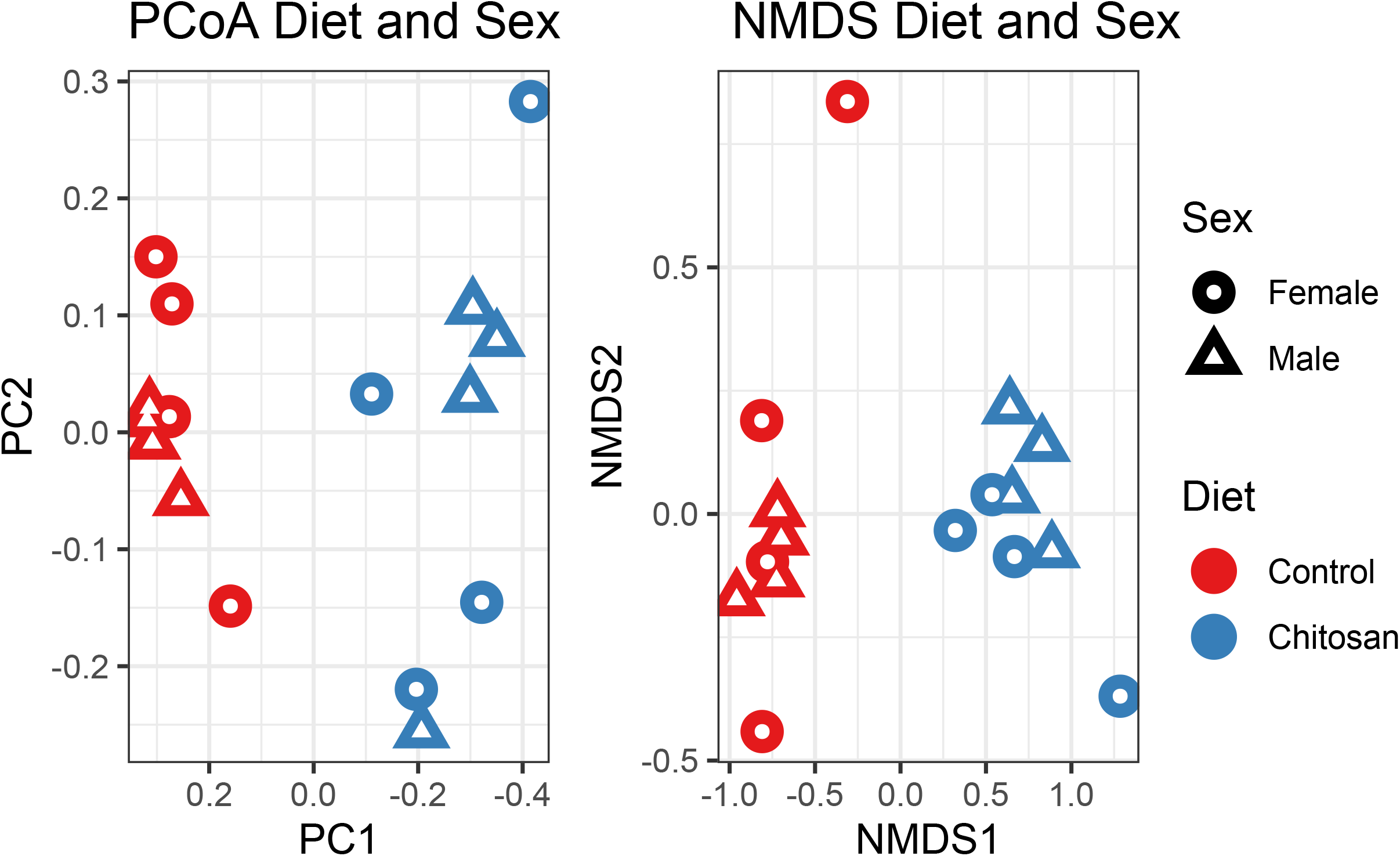
Ordination of house fly microbial community relative abundances calculated using (A) Unifrac distances for principal coordinates analysis (PCoA) and (B) Bray Curtis distances for non-metric multidimensional scaling (NMDS). Symbols denote fly microbial communities, with red representing chitosan diet and blue representing control diet flies (Diet PERMANOVA, F= 28.831, P<0.001). Circles are females, and triangles are males, though there was no significant difference in community composition by sex (Sex PERMANOVA, F=1.084, P<0.274 n.s.).

### 3.3 Chitosan increases bacterial load fly microbiomes

We targeted the V4 region of the 16S rRNA gene and used qPCR to quantify the differences in the number of gene copies between the two diets. We referenced the original fly sample weights before DNA extraction to enumerate the gene copy number per fly and the gene copy number per unit biomass. On average, the chitosan-fed flies weighed 26% less than flies fed a control diet (t-test, P<0.05, **Figure 3A**). Contrary to our expectations, chitosan-fed flies had 56% more the bacteria in their microbiome as measured by 16S rRNA gene copy number per fly (t-test, P<0.001, **Figure 3B**) and 2.02 times greater copy number per unit fly biomass (t-test, P=0.08, **Figure 3C**).

**Figure 3.**
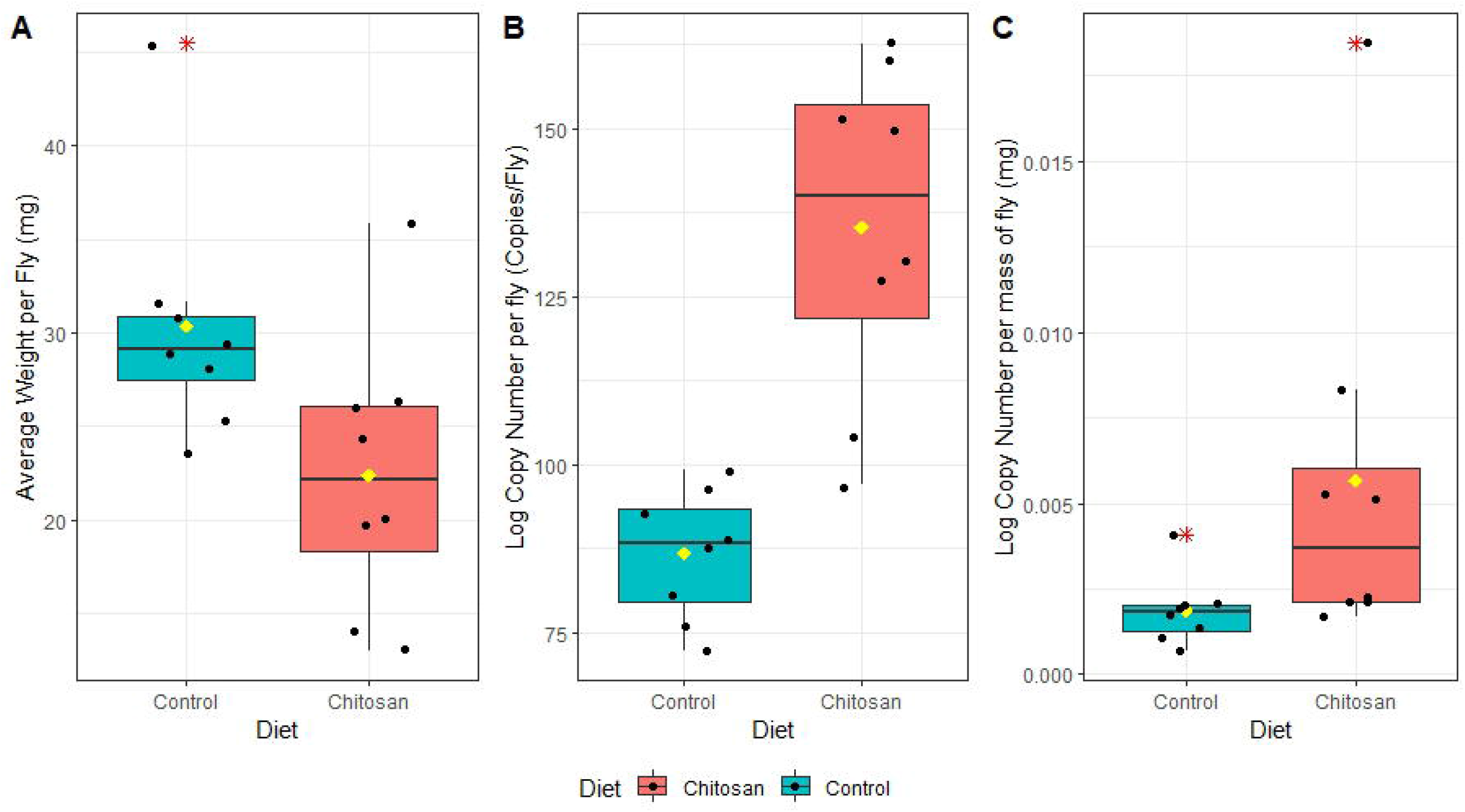
Diet had a significant effect on body weight and fly microbiome bacterial load. Boxplots show the median, interquartile range, and 95% confidence intervals for (A) Average weight per fly (mean biomass/fly =30.4 and 22.4 mg for control and chitosan, respectively). (B) Log copy number per fly (mean log copies/fly = 86.7 and 135.3 for control and chitosan, respectively). (C) Log copy number per mass of fly (mean log copies/mg = 1.87E-3 and 5.67E-3 for control and chitosan, respectively). The black points show values for individual samples. Red asterisk shows data outliers. Yellow point shows the mean.

### 3.4 Chitosan alters gut microbial community structure

To gain a perspective on how a chitosan diet affects taxonomic composition, we examined the relative abundances of the top 5 groups of each taxonomic rank for class, family, and genus (**Figure 4A**). The chitosan panel shows eight chitosan diet replicates (n=4 female, and n=4 male). *Gammaproteobacteria* is the most abundant class in all eight trials, with over 80% relative abundance of this group in all eight trials. *Burkholderiaceae* is the most abundant family in all the control diet flies, with over 70% relative abundance of this group (**Figure 4B**). Conversely, *Yersiniaceae* is the most abundant family in chitosan-fed diet flies, with over 40% relative abundance of this group (**Figure 4B**). *Ralstonia* is the most abundant genus in control diet flies, with over 40% relative abundance of this group in all trials except CF2 (**Figure 4C**). In chitosan fed flies, *Ralstonia* are present but *Serratia* is the most abundant genus in all eight trials, with over 30% relative abundance of this group (**Figure 4C**). In analyzing ASVs by genus, we observed an average relative abundance in control flies of *Kosakonia:* 0%, *Providencia:* 10.41%, *Ralstonia:* 57.92%, *Serratia:* 0.04%; and in chitosan-fed flies of *Kosakonia:* 4.04%, *Providencia:* 1.30%, *Ralstonia:* 20.30%, *Serratia*: 59.89%.

**Figure 4.**
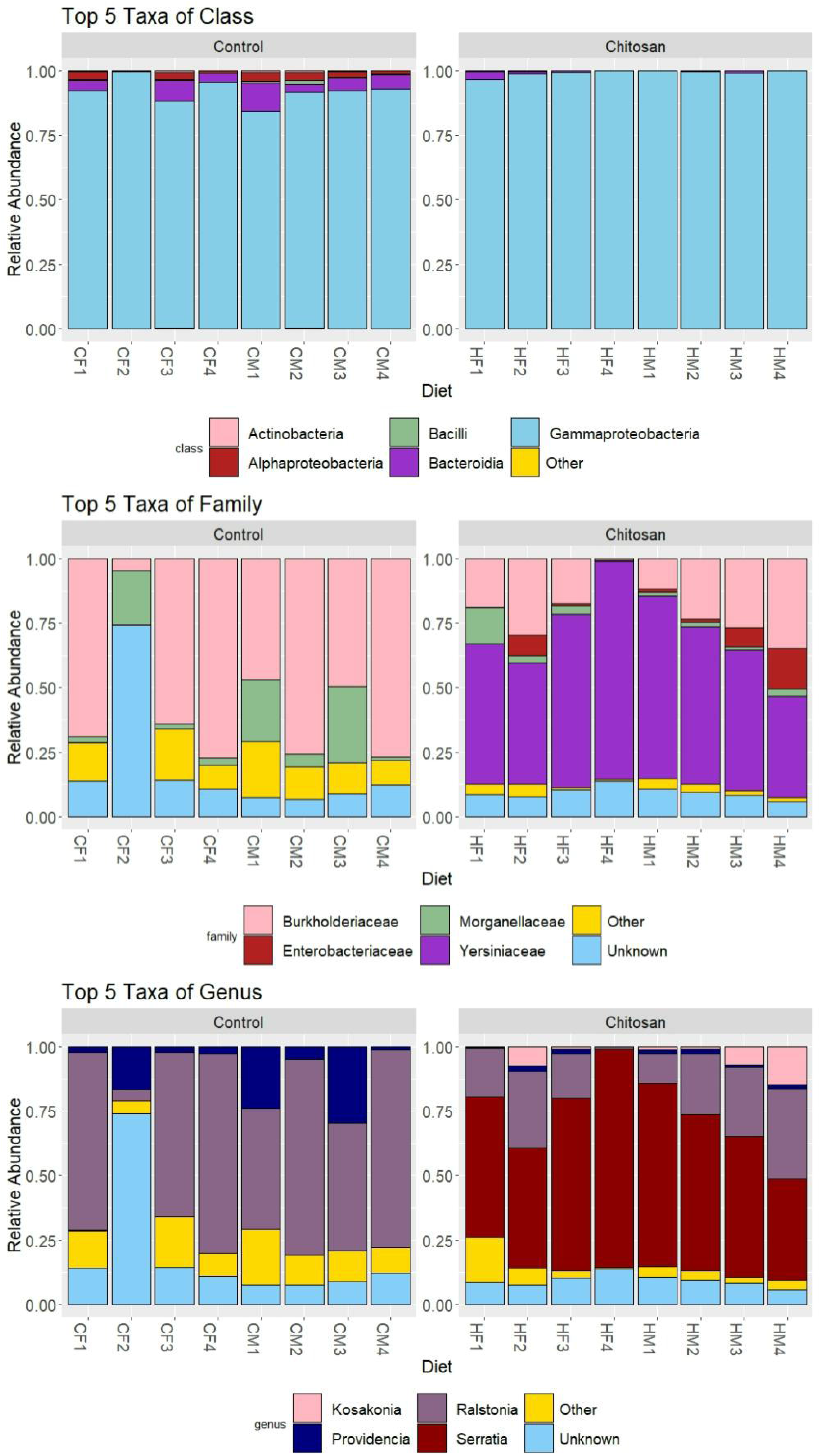
Taxonomic composition varies between samples and diet treatments. (A) Relative abundances of top 5 class groups in house flies. (B) Relative abundances of top 5 family groups in house flies. (C) Relative abundances of top 5 genus groups in house flies. In each display, the control panel shows eight control diet replicates, female, (CF1-CF4) and male (CM1-CM4).

We suspect that the enrichment of *Serratia* spp. could be due to their ability to efficiently degrade chitosan compared to other bacteria (Vaaje-Kolstad et al. 2013). *Serratia* was absent in all control groups except in one control female trial where *Serratia* was less than 1% relative abundance. In our study, *Serratia* is not a well-established member of the fly gut microbiome on a sucrose diet. When chitosan is added to the diet, *Serratia* becomes a dominant group in the fly gut microbiome. Though total bacteria increase in chitosan-fed flies, the abundance of some dominant taxa, including *Ralstonia* species, decreases. While there are many studies on the antifungal effects of a chitosan solution diet, there are few studies featuring antibacterial effects of a chitosan solution diet. Evidence suggesting that chitosan is an effective antibacterial agent includes a study conducted by Li et al. (2010), where varying concentrations of chitosan in growth media negatively affects *Serratia marcescens* strains’ ability to grow. After 30 min of incubation in 0.10 mg chitosan/mL, both strains were unable to grow (Li et al. 2010). Moye et al. (2014) shows chitosan solution at different concentrations effectively kills *Serratia*. It’s possible a 2% chitosan fly diet is not toxic to *Serratia*, but at higher concentrations its toxicity reduces the abundance of *Serratia*. Because chitosan is a known antifungal, and fungal species are detected in wild (but not lab-cultivated) house fly microbiome (Park et al., 2017), this aspect of chitosan mode of action should be examined in future studies.

Chitosan increases house fly mortality (Stoffolano et al. 2020), but the mechanism behind chitosan’s effect in killing adult flies may be through deleterious effects on the gut microbiome of house flies. There are various suggested modes of actions of chitosan on bacteria. One theory suggests that chitosan binds to the outer membrane of gram-negative bacteria and blocks the flow of nutrients therefore causing DNA damage (Kravanja et al. 2019). Some degradation products of chitosan, such as N-acetylglucosamine, can be used as a carbon source by compatible bacteria, such as *Serratia* spp. and *Ralstonia* spp. Degradation products of chitosan that are antimicrobial, like chitosan-oligosaccharides (COS), could be produced in high concentrations when chitosan is supplemented in a diet. In this way, the efficient degradation of chitosan by some community members would negatively affect susceptible bacteria in the community. COS can bind to negatively charged O-antigen of LPS on bacterial cell walls, which blocks nutrient flow (Kravanja et al. 2019). We hypothesize that chitosan is degraded by chitinolytic bacteria, which increases the number of antimicrobial peptides like COS in the environment. The blocking of nutrient flow on bacterial cell walls within the fly gut caused by the COS antimicrobial peptides (Kravanja et al. 2019) may be a plausible explanation for high mortality rates of flies on a chitosan supplemented diet.

Recent studies have been exploring the insecticidal effectiveness of sugar baits such as xylitol and erythritol. Alone, xylitol would reduce sample population to 50% by day 3 and erythritol by day 2. However, when the sugar baits would be mixed in with sucrose, the insecticidal effectiveness had a life-extending effect. There are suggestions leading towards fly dehydration or caloric starvation that could be affecting longevity, however there are other literature that suggest that these sugar alcohol diets decrease the bacterial communities due to their antimicrobial properties or effects of biofilm development (Uhari et al., 2000; de Cock et al., 2016; Burgess et al., 2018; Burgess et al., 2021). More research needs to be done to determine the mode of action of carbohydrates that affect fly longevity. However, our study shows that chitosan diet alters community structure and dramatically increases bacterial load while also decreasing fly weight, pointing to a mechanism of disrupted host nutrient uptake function.

### 3.5 Chitosan decreases gut richness and diversity

We further hypothesized that chitosan would reduce gut microbiome richness, and this reduced microbiome diversity would be associated with fly mortality (Gould et al., 2018). Chitosan diet reduced richness but not evenness of fly microbiomes. We analyzed the alpha diversity metrics between the control and chitosan fed fly microbiome. We performed t-tests to ask if these diversity metrics varied between diets. The observed richness of the control group was 32.13, while the chitosan group had an observed richness of 16.00 (t-test P<0.001, **Figure 5A**). Pielou’s evenness was 0.51 for the control and 0.46 for chitosan, indicating highly uneven committees in both groups, with no significant difference in evenness between the two diets (**Figure 5B**). Shannon diversity was 1.75 for the control and 1.28 for the chitosan bacterial communities (t-test P<0.05, **Figure 5C**).

**Figure 5.**
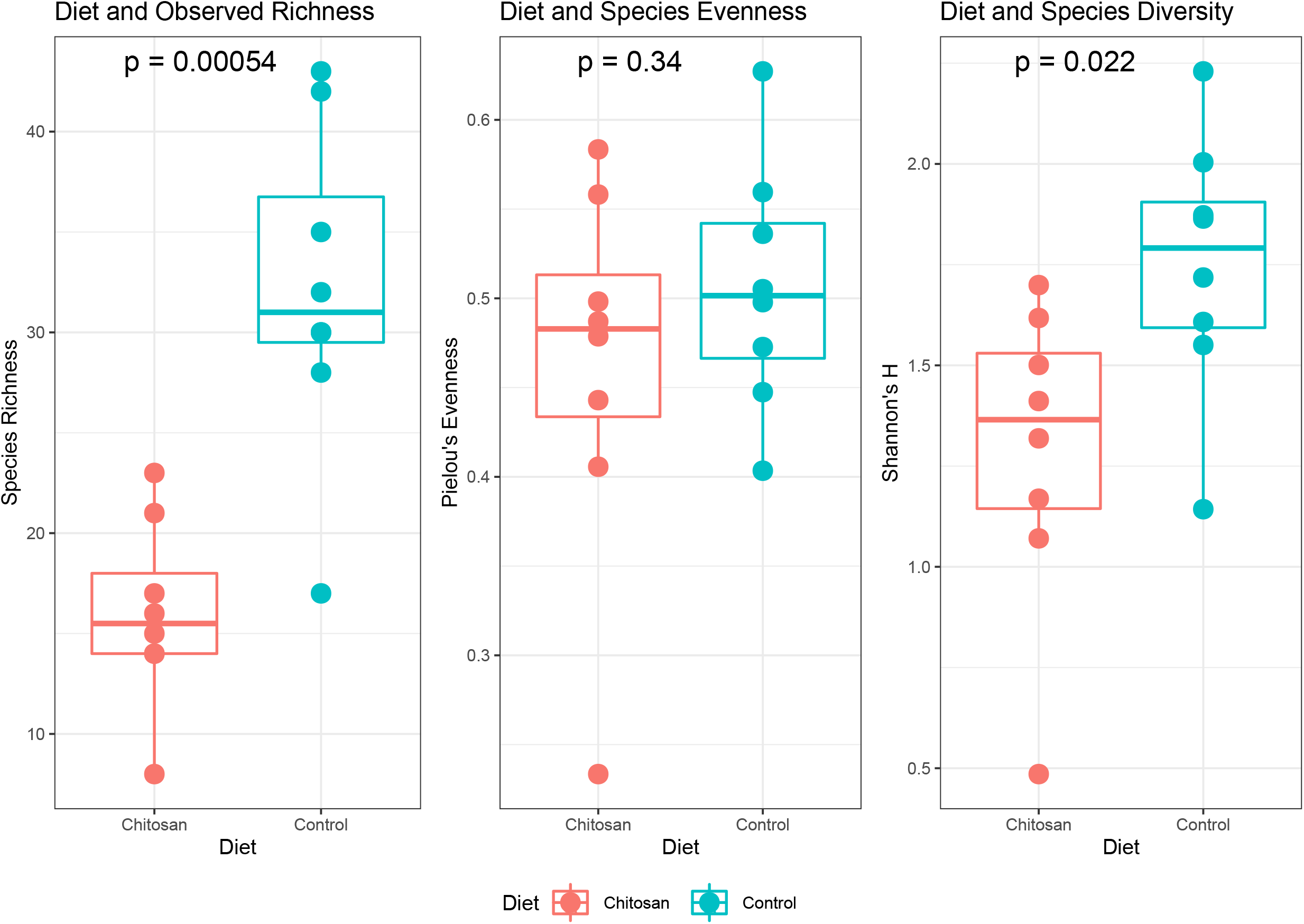
Boxplots show observations for each diversity metric. Species richness is the total number of observed bacterial species (A). To evaluate community evenness, we used Pielou’s evenness metric. Species evenness accounts for the distribution of abundances for species in a community. Values range from 0 to 1, where higher values indicate greater community evenness. (B) To evaluate community diversity, we used Shannon’s H metric. Shannon diversity accounts for both richness and evenness (C).

Our data shows that bacteria community richness and diversity significantly decrease with the addition of 2% chitosan in the house fly diets compared to the control diet. As expected, the community evenness for both diets were low. This is likely because both groups lived in a regulated laboratory environment where there was not much exposure to the bacterial biodiversity a typical wild fly would encounter. A shift in bacterial abundance has been previously observed between fly habitat and location (Park et al. 2019), where wild flies differ from lab flies’ bacterial abundance. In our case, the environment was consistent between treatments, with the only difference being the introduction of chitosan to one group.

Even though chitosan fed flies weighed less, they had more bacteria based on gene copy number per fly and gene copy number per mass of fly. Chitosan decreases richness and diversity in the gut microbiome of houseflies, but chitosan does not lower the bacterial abundance in the gut microbiome of house flies. We reject our hypothesis that chitosan decreases bacterial abundance.

A possible explanation for the decreased community richness and diversity in the chitosan diet would be that chitosan negatively impacts bacteria in the gut microbiome, therefore lowering the total number of bacterial species. Chitosan may also affect specific bacteria negatively, therefore lowering some, but not all, bacterial taxa in the gut microbiome. It is also possible that chitosan-amended diets encourage the growth of bacteria that can use chitosan as a food source, allowing them to out-compete the natural gut flora.

## 4 Conclusion and Future Directions

Our results confirm that adding chitosan to house fly diets increases their mortality, and this is accompanied by stark changes in microbial community composition and abundance. Chitosan affects the microbiome of house flies, but not in the way we had hypothesized. Chitosan is proven to be an antifungal agent, and our research suggests the possibility that chitosan shows selective antibacterial activity, with an overall effect of increasing bacterial abundance in the fly microbiome. With this new information, we can better utilize chitosan to target specific pathogens. The mode of chitosan’s antibacterial effect is still unverified. This study gives a piece of the puzzle, demonstrating how specific gut bacteria respond to chitosan.

## Conflict of Interest

*The authors declare that the research was conducted in the absence of any commercial or financial relationships that could be construed as a potential conflict of interest*.

## Author Contributions

**Hila Schaal:** Conceptualization, Data curation, Formal analysis, Investigation, Methodology, Software, Validation, Visualization, Writing-Original draft preparation. **Mallory Choudoir:** Data curation, Methodology, Writing-Reviewing and Editing. **Vedang Diwanji:** Data curation, Investigation. **John Stoffolano Jr.:** Conceptualization, Funding acquisition, Resources, Writing-Reviewing and Editing. **Kristen DeAngelis:** Conceptualization, Funding acquisition, Project administration, Resources, Supervision, Writing-Reviewing and Editing.

## Funding

The major part of this work was supported by funding from the National Institute of Food and Agriculture, U.S. Department of Agriculture, the Massachusetts Agricultural Experiment Station, and the Department of Microbiology. The work was also partially supported by USDA/NIFA support to the Stockbridge School of Agriculture at the University of Massachusetts, Amherst, under Hatch project MAS00527 to JGS. The contents are solely the responsibility of the authors and do not necessarily represent the official views of the USDA or NIFA.

## Acknowledgments

We would like to acknowledge the contributions of the following colleagues and institutions: Dr. Christopher Geden for fly pupae supply, Dr. John Burand for various laboratory equipment, Dr. Samuel Hazen for gel equipment, Dr. Ravi Ranjan and the Genomics Resource Lab at University of Massachusetts Amherst for Illumina sequencing and quality control. Thanks also to Eitan Altabet for help with fly growth, dissections, and sample preparation.

## Data Availability Statement

The sequence data for this study can be found in the GenBank database under accession numbers OL416221 - OL416327.

## Notes

### Competing Interest Statement

The authors have declared no competing interest.

## References

Andongma, A.A., Wan, L., Dong, X.-P., Akami, M., He, J., Clarke, A.R., and Niu, C.-Y. (2018). The impact of nutritional quality and gut bacteria on the fitness of *Bactrocera minax* (Diptera: Tephritidae). R. Soc. Open. Sci. 5, 180237.

Apprill, A., McNally, S., Parsons, R., & Weber, L. (2015). Minor revision to V4 region SSU rRNA 806R gene primer greatly increases detection of SAR11 bacterioplankton. Aquatic Microbial Ecology, 75(2), 129–137. https://doi.org/10.3354/ame01753

Bahrndorff, S., Jonge, N. de, Skovgård, H., & Nielsen, J. L. (2017). Bacterial Communities Associated with Houseflies *(Musca domestica* L.) Sampled within and between Farms. PLOS ONE, 12(1), e0169753. https://doi.org/10.1371/journal.pone.0169753

Behar, A., Yuval, B., & Jurkevitch, E. (2008). Gut bacterial communities in the Mediterranean fruit fly *(Ceratitis capitata)* and their impact on host longevity. Journal of Insect Physiology, 54(9), 1377–1383. https://doi.org/10.1016/j.jinsphys.2008.07.011

Ben-Yosef, M., Behar, A., Jurkevitch, E., Yuval, B. (2008). Bacteria-diet interactions affect longevity in the medfly-*Ceratitis capitata*. J. Appl. Entomol. 132(9□10):690 – 694

Brewer, N., McKenzie, M. S., Melkonjan, N., Zaky, M., Vik, R., Stoffolano, J. G., & Webley, W. C. (2021). Persistence and Significance of Chlamydia trachomatis in the Housefly, *Musca domestica L*. Vector Borne and Zoonotic Diseases (Larchmont, N.Y.), 21(11), 854–863. https://doi.org/10.1089/vbz.2021.0021

Burgess IV, Edwin R., E. E. Taylor, Anthony Acevedo, Michaela Tworek, Dana Nayduch, Neetika Khurana, Jon S. Miller, and Christopher J. Geden. “Diets of erythritol, xylitol, and sucrose affect the digestive activity and gut bacterial community in adult house flies.” Entomologia Experimentalis et Applicata 169, no. 10 (2021): 878–887.

Burgess IV, Edwin R., and B. H. King. “Insecticidal potential of two sugar alcohols to Musca domestica (Diptera: Muscidae).” Journal of economic entomology 110, no. 5 (2017): 2252–2258.

Callahan, B. J., McMurdie, P. J., Rosen, M. J., Han, A. W., Johnson, A. J. A., & Holmes, S. P. (2016). DADA2: High-resolution sample inference from Illumina amplicon data. Nature Methods, 13(7), 581–583. https://doi.org/10.1038/nmeth.3869

Caporaso, J. G., Ackermann, G., Apprill, A., Bauer, M., Berg-Lyons, D., Betley, J., Fierer, N., Fraser, L., Fuhrman, J. A., Gilbert, J. A., Gormley, N., Humphrey, G., Huntley, J., Jansson, J. K., Knight, R., Lauber, C. L., Lozupone, C. A., McNally, S., Needham, D. M., … Weber, L. (2018, April 14). EMP 16S Illumina Amplicon Protocol. Protocols.Io. https://www.protocols.io/view/emp-16s-illumina-amplicon-protocol-nuudeww

Caporaso, J. G., Lauber, C. L., Walters, W. A., Berg-Lyons, D., Huntley, J., Fierer, N., Owens, S. M., Betley, J., Fraser, L., Bauer, M., Gormley, N., Gilbert, J. A., Smith, G., & Knight, R. (2012). Ultra-high-throughput microbial community analysis on the Illumina HiSeq and MiSeq platforms. The ISME Journal, 6(8), 1621–1624. https://doi.org/10.1038/ismej.2012.8

de Cock, Peter, Kauko Mäkinen, Eino Honkala, Mare Saag, Elke Kennepohl, and Alex Eapen. “Erythritol is more effective than xylitol and sorbitol in managing oral health endpoints.” International journal of dentistry 2016 (2016).

de Jonge, N., Michaelsen, T. Y., Ejbye-Ernst, R., Jensen, A., Nielsen, M. E., Bahrndorff, S., & Nielsen, J. L. (2020). Housefly *(Musca domestica L.)* associated microbiota across different life stages. Scientific Reports, 10(1), 7842. https://doi.org/10.1038/s41598-020-64704-y

Engel, P., and Moran, N.A. (2013). The gut microbiota of insects - diversity in structure and function. FEMS Microbiol. Rev. 37, 699–735.

Geden, C. J., Nayduch, D., Scott, J. G., Burgess IV, E. R., Gerry, A. C., Kaufman, P. E., Thomson, J., Pickens, V. and Machtinger, E. T. 2021. House fly (Musca domestica [Dip;tera: Muscidae]) – Biology, pest status, current management prospects, and research needs. Jour. Integrated Pest Management. 12: 1–38.

Gould, Alison L., Vivian Zhang, Lisa Lamberti, Eric W. Jones, Benjamin Obadia, Nikolaos Korasidis, Alex Gavryushkin, Jean M. Carlson, Niko Beerenwinkel, and William B. Ludington. “Microbiome interactions shape host fitness.” Proceedings of the National Academy of Sciences 115, no. 51 (2018): E11951–E11960.

Gupta, A. K., Nayduch, D., Verma, P., Shah, B., Ghate, H. V., Patole, M. S., & Shouche, Y. S. (2012). Phylogenetic characterization of bacteria in the gut of house flies (Musca domestica L.). FEMS Microbiology Ecology, 79(3), 581–593. https://doi.org/10.1111/j.1574-6941.2011.01248.x

Hahn, T., Tafi, E., Paul, A., Salvia, R., Falabella, P., & Zibek, S. (2020). Current state of chitin purification and chitosan production from insects. Journal of Chemical Technology & Biotechnology, 95(11), 2775–2795. https://doi.org/10.1002/jctb.6533

Kong, M., Chen, X. G., Xing, K., & Park, H. J. (2010). Antimicrobial properties of chitosan and mode of action: A state of the art review. International Journal of Food Microbiology, 144(1), 51–63. https://doi.org/10.1016/j.ijfoodmicro.2010.09.012

Kralik, P., & Ricchi, M. (2017). A Basic Guide to Real Time PCR in Microbial Diagnostics: Definitions, Parameters, and Everything. Frontiers in Microbiology, 8. https://doi.org/10.3389/fmicb.2017.00108

Kravanja, G., Primožič, M., Knez, Ž., & Leitgeb, M. (2019). Chitosan-Based (Nano)Materials for Novel Biomedical Applications. Molecules, 24(10), 1960. https://doi.org/10.3390/molecules24101960

Li, B., Su, T., Chen, X., Liu, B., Zhu, B., Fang, Y., Qiu, W., & Xie, G. (2010). Effect of chitosan solution on the bacterial septicemia disease of Bombyx mori (Lepidoptera: Bombycidae) caused by *Serratia marcescens*. Applied Entomology and Zoology, 45(1), 145–152. https://doi.org/10.1303/aez.2010.145

Martin, M. (2011). Cutadapt removes adapter sequences from high-throughput sequencing reads. EMBnet. Journal, 17(1), 10–12. https://doi.org/10.14806/ej.17.1.200

McMurdie, Paul J., and Susan Holmes. “phyloseq: an R package for reproducible interactive analysis and graphics of microbiome census data.” PloS one 8, no. 4 (2013): e61217.

McLaren, M. R. (2020). Silva SSU taxonomic training data formatted for DADA2 (Silva version 138) [Data set]. Zenodo. https://doi.org/10.5281/zenodo.3986799

Moye, Z. D., Burne, R. A., & Zeng, L. (2014). Uptake and Metabolism of N-Acetylglucosamine and Glucosamine by *Streptococcus* mutans. Applied and Environmental Microbiology, 80(16), 5053–5067. https://doi.org/10.1128/AEM.00820-14

Oksanen, Jari, Roeland Kindt, Pierre Legendre, Bob O’Hara, M. Henry H. Stevens, Maintainer Jari Oksanen, and M. A. S. S. Suggests. “The vegan package.” Community ecology package 10, no. 631-637 (2007): 719.

Orzali, L., Corsi, B., Forni, C., & Riccioni, L. (2017). Chitosan in Agriculture: A New Challenge for Managing Plant Disease. https://doi.org/10.5772/66840

Pan, H., Fu, C., Huang, L., Jiang, Y., Deng, X., Guo, J., & Su, Z. (2018). Anti-Obesity Effect of Chitosan Oligosaccharide Capsules (COSCs) in Obese Rats by Ameliorating Leptin Resistance and Adipogenesis. Marine Drugs, 16(6). https://doi.org/10.3390/md16060198

Parada, A. E., Needham, D. M., & Fuhrman, J. A. (2016). Every base matters: Assessing small subunit rRNA primers for marine microbiomes with mock communities, time series and global field samples. Environmental Microbiology, 18(5), 1403–1414. https://doi.org/10.1111/1462-2920.13023

Park, R., Dzialo, M. C., Spaepen, S., Nsabimana, D., Gielens, K., Devriese, H., Crauwels, S., Tito, R. Y., Raes, J., Lievens, B., & Verstrepen, K. J. (2019). Microbial communities of the house fly *Musca domestica* vary with geographical location and habitat. Microbiome, 7(1), 147. https://doi.org/10.1186/s40168-019-0748-9

Pickens, L.G., and Miller, R.W. Techniques for trapping flies on dairy farms. (1987). J. Agric. Entomol. (4)4: 305–313

Pinero, Jaime & Chiu, Katherine & Colletti, Kay & Dixon, Zoe & Salemme, Victoria & Crnjar, Roberto & Sollai, Giorgia. (2021). Effects of chitosan and erythritol on labellar taste neuron activity, proboscis extension reflex, daily food intake, and mortality of male and female spotted-winged drosophila, *Drosophila suzukii*. Journal of insect physiology. 131. 104240. 10.1016/j.jinsphys.2021.104240.

R Core Team. “R: A language and environment for statistical computing.” (2013).

Rabea, E. I., Badawy, M. E.-T., Stevens, C. V., Smagghe, G., & Steurbaut, W. (2003). Chitosan as Antimicrobial Agent: Applications and Mode of Action. Biomacromolecules, 4(6), 1457–1465. https://doi.org/10.1021/bm034130m

Rabea, E. I., Nasr, H. M., Badawy, M. E. I., & El-Gendy, I. R. (2015). Toxicity of naturally occurring Bio-fly and chitosan compounds to control the Mediterranean fruit fly *Ceratitis capitata* (Wiedemann). Natural Product Research, 29(5), 460–465. https://doi.org/10.1080/14786419.2014.948873

Stoffolano, J. G., Jr. 2019. Fly foregut and transmission of microbes. Adv. Insect Physiol. 57: 29–95.

Stoffolano, J., Wong, R., Lo, T., Ford, B., & Geden, C. J. (2020). Effect of chitosan on adult longevity when fed, in no-choice experiments, to *Musca domestica* L., *Tabanus nigrovittatus* Macquart, and *Phormia regina* (Meigen) adults and its consumption in adult *Musca domestica L*. Pest Management Science, 76(12), 4293–4300._https://doi.org/10.1002/ps.5996

Telmadarrehei, T., Tang, J. D. Raji, O. Rezazadeh, A. Narayanan, L. Shmulsky, R and Jeremic. D. 2020. A Study of the gut bacterial community of *Reticulitermes virginicus* exposed to chitosan treatment. Insects 11: 681. https://doi.org/10.3390/insects11100681

Uhari, Matti, Terhi Tapiainen, and Tero Kontiokari. “Xylitol in preventing acute otitis media.” Vaccine 19 (2000): S144–S147.

Vaaje-Kolstad, G., Horn, S. J., Sørlie, M., & Eijsink, V. G. H. (2013). The chitinolytic machinery of *Serratia marcescens* – a model system for enzymatic degradation of recalcitrant polysaccharides. The FEBS Journal, 280(13), 3028–3049. https://doi.org/10.1111/febs.12181

